# Tree ring segmentation performance in highly disturbed trees using deep learning

**DOI:** 10.1101/2025.03.14.643209

**Authors:** Joe David Zambrano-Suárez, Jorge Pérez-Martín, Alberto Muñoz-Torrero Manchado, Juan Antonio Ballesteros Cánovas

## Abstract

Dendrogeomorphology has provided valuable insights for dating geomorphic events, but requires the challenging analyses of tree-ring records from highly disturbed trees. Deep learning algorithms have been successfully used to detect ring boundaries under normal tree growth conditions. Here, we test if deep learning can perform tree ring segmentation in highly abnormal growth patterns. To this end, this study explores the relation between the complexity of convolutional neural networks (CNN)-based architectures, cellular detail levels, and the capacity to segment ring borders in abnormal tissues. Increment cores were collected from a debris flow-affected area in the Pyrenees, while images were acquired using a digital camera with a high-resolution macro. We defined four sets of experiments, including varying image resolution through downsampling, applying different architectures, and using image filters. Moreover, we test if the inclusion of the growth direction into a patchify-based system applied to increment cores improves the performance of the system. Our results suggest that intelligent systems can recognize tree-rings boundaries, but their performance was lower with high abnormal growth patterns due to the significant differences in colors and textures from normal growth patterns. However, the proposed segmentation system was able to segment sets of narrow ring borders, spaced above 200 μm, where the color remained unchanged. Interestingly, our results suggest that the model ignored cellular details and relied on color gradients to detect ring borders when analyzing at the macro level. This implies that the image resolution is only becoming critical for densely packed rings with minimal spacing. Finally, we observed that CNN-based segmentation systems were unable to infer growth direction based solely on tree ring convexity and cellular details within an increment core patch. Our results provide new insights into how deep learning could be used in tree-ring research, but they still reveal the existing challenges with disturbed trees.

## Introduction

Dendrochronology is an absolute dating method that uses the structure of tree rings to determine the age of trees and analyze past climate conditions [1], making it a valuable tool in scientific fields such as climatology, ecology, archaeology, hydrology, and geomorphology, among others [2–5]. In particular, the use of tree rings to determine the spatial and temporal patterns of geomorphic processes is known as dendrogeomorphology [6]. Extreme events such as floods, debris flows, landslides, or snow avalanches frequently affect existing vegetation, disrupting the normal growth patterns of trees and shrubs [4,5,7,8]. As a result, affected trees often exhibit growth disturbances characterized by unusual ring patterns, such as changes in growth rates, variations in tissue coloration, or differences in texture [9]. Dating and identifying these growth disturbances is challenging and heavily reliant on expert analysis.

Classical tree-ring dating methods involve the use of binoculars attached to a measuring table operated by an expert [10]. This optical approach is highly accurate, but requires significant time while discrepancies in measurements between experts’ interpretations may appear [11]. Thus, optical measurements often rely on mechanical-assisted techniques to assist in marking, outlining, and documenting the boundaries and widths of the tree rings [12–14]. Recently, the employ of digital images and microelectronics technology has become more popular due to their efficiency and cost-benefits ratios, with microelectronics measurements obtaining tree ring images from a highly accurate scanners that are then digitally analyzed using specific computer software [14–18]. The digital analyses of samples may introduce semi-automatic algorithms to detect ring boundaries based on changes in pixel intensities [15,16,18], but it needs to be assisted by an expert to correct measurements and misinterpretation.

Deep Learning (DL) and Convolutional Neural Networks (CNNs) have been used successfully across many fields for identifying visual patterns and making predictions [19–21]. CNNs extract complex information about the context of the pixels and work with complex and abstract information [22]. In semantic segmentation, where the objective is pixel-level classification, the encoder-decoder architecture is one of the most widely adopted approaches [23,24]. This includes notable structures such as U-Net, V-Net, Deeplab and Fully Convolutional Network (FCN) [25–28] . U-Net demonstrates remarkable capabilities in image analysis [29], providing highly accurate image descriptions and serving as the foundation for numerous state-of-the-art DL architectures specifically developed and trained for image segmentation tasks [30,31]. In recent years, CNN-based systems have been employed in the task of segmenting growth tree rings for dendrochronology [10,11,14,32–38]. García-Hidalgo et al. [38] designed a system that achieves promising results in automatic segmentation, but specifically designed to work with wood-anatomical images from thin microsections, making it unsuitable for studies involving full increment tree cores. Gillert et al. [33] introduces Iterative Next Boundary Detection (INBD), a system for cross-section microscopy images, which was later studied on tree cross-section images at the macroscopic level [32]. Several studies have therefore focused on macroscopic images obtained by increment tree cores [14,36,37], even using full cross sections [34,35] although training and performing image segmentation using rectangular images as increment cores. These macroscopic-based studies reveal dissimilar performance, highlighting the need for further research.

In the literature, systems have been developed to successfully segment growth tree ring boundaries in easily recognized and normal growth patterns from increment cores. However, a knowledge gap exists in the successful segmentation of tree ring boundaries from trees presenting abnormal growth patterns or densely packed tree ring sets at a macroscopic level [32,34–37]. Moreover, systems with varying levels of cellular detail have shown robust performance, but the interaction between architecture complexity and cellular detail, and its impact on segmentation effectiveness, remains unexplored. To the best of our knowledge, only two studies have analyzed these interactions [32,35]. Specifically, Marichal & Randall [32] successfully implemented a system originally designed for microscopic cross sections to macro cross sections [33], although the method requires the cross sections to be noise-free. Also, Kim et al. [35] has experimented with different levels of cellular detail, scanner and mobile camera, concluding that higher cellular detail yields improved results due to the internal functionality of the CNN-based architecture employed. However, their study exclusively focuses on well-defined tree ring boundaries, cautiously avoiding irregular tree rings, piths, and other noise sources such as mechanical disturbances. Lastly, the question of whether systems using increment tree cores images with cellular detail at a macroscopy level can operate effectively without incorporating prior knowledge of tissue growth direction remains still unclear.

Here, we focus on the use of systems to distinguish tree-ring growth patterns in highly disturbed trees. To do that, we perform different experiments with increment cores with growth disturbances caused by past mass-movement processes including: (i) different architectures of varying complexity (ii) diverse image preprocessing techniques. By analyzing the resulting segmentation, both quantitative and qualitative, we aim to address several key research questions (RQ):

− RQ: How does the complexity of U-Net architecture impact performance of system in the segmentation of tree ring boundaries?
− RQ: What is the influence of the detailed cellular level of increment tree cores on the effectiveness of system in the segmentation of tree ring boundaries?
− RQ: Does the performance of the system change when images are not adjusted to ensure a consistent growth direction?
− RQ: Does the performance of a system for segmenting tree ring boundaries in abnormal growth patterns improve when more samples with growth disturbances are included in the dataset?

Through these research questions, we seek to gain a deeper understanding of how the segmentation of tree ring borders in growth patterns affected by growth disturbance is related to architecture and image complexity.

## Materials and Methods

### Data collection

We used increment bores to extract wood samples from *Pinus sylvestris* L. individuals growing in a debris flow cone at Pineta Valley (Spanish Pyrenees) (S1 Appendix). Sampled trees presented external evidence of past debris flow, such as decapitated, tilted, scarred or buried trees [7]. This implies that samples include anomalous growth patterns such as abrupt suppression/releases, reaction wood and callus tissues [9] (Fig 1). After field collection, samples were sanded and polished using sandpaper (from 120 to 800 grit) to enhance tree ring visibility. Then, all samples were digitized using the CaptuRING tool [39]. This device was equipped with a Digital Camera EOS R8 with a macro EF100mm lens, specifically designed to capture fine textures and details in close-ups. The system was complemented by an MF18 Macro Flash, providing uniform and controlled illumination. A stitching process on the sequential images was later performed using an Image Composite Editor (ICE) software (Microsoft Research ^TM^). Additionally, three extra images were reserved as one of the test datasets, but these were photographed under lamps emitting a warm yellowish light rather than using the Macro Flash.

**Fig 1.**
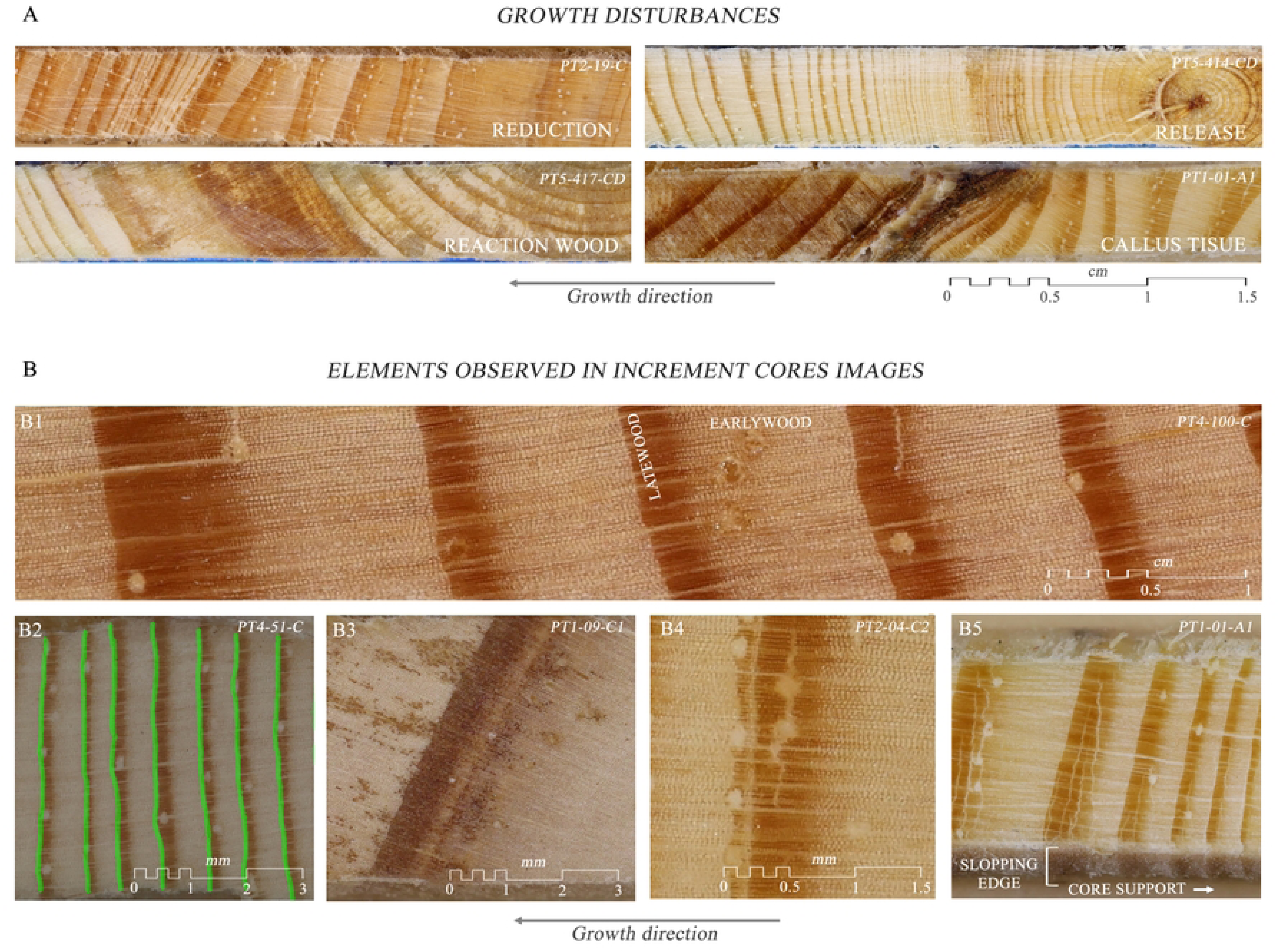
Summary of Information in increment core images. (A) Summary of the growth disturbances presented in the images of increment cores. Including growth realease, growth reduction, callus tissue and reaction wood, (B) Summary of elements present in the images of increment cores. (B1) Transitions from earlywood to latewood. (B2) Typical case of a ring border. (B3) Case not labeled as a ring border due to no reduction in cell size. (B4) Case not labeled as a ring border due to a junction at a specific point between two rings. (B5) Case of the presence of sloping edges with ring borders and regions with core support.

Overall, images from 64 increment tree cores have been obtained, showing a wide range of growth tree rings (Table 1) and a total number of 11,947 tree rings. Cells in earlywood and their borders were easily distinguishable due to their larger lumen sizes. In contrast, latewood cells were only identifiable in certain regions and primarily during their early stages, becoming indistinct in later stages (Fig 1). Consequently, we used a Pixel Annotation Tool [40] in a digital drawing tablet to label the transition between latewood and earlywood as tree ring border. One of the main challenges lay in ensuring consistency in the labeling of false tree rings, as their distinction from standard tree rings could vary in degree of similarity. In our dataset, tree ring borders were labeled as those transitions characterized by light-colored bands and larger cells following a late latewood stage with dark and small cells (Fig 1). We did not label a tree ring border when there was a transition in coloration, but the cells still maintained a considerable relative size (Fig 1). Also, when there were two transitions from earlywood to latewood in close proximity, with no earlywood in between in one section, the first suspected tree ring border was not labeled as a border (Fig 1). Additionally, the images not only included core tissues captured perpendicularly to view of the field of the camera but also featured areas with sloping edges, some containing tree ring borders that were not labeled as such, as well as regions of the core support (Fig 1).

**Table 1.** Summary characteristics of the increment core images.

| Quantile | Width (px) | Height (px) | Pixels labeled (%) | Number of tree rings |
| --- | --- | --- | --- | --- |
| 0 | 6,588 | 502 | 2.52 | 47 |
| 0.25 | 25,821.75 | 789.75 | 5.43 | 106 |
| 0.5 | 32,350.5 | 853.5 | 7.34 | 169 |
| 0.75 | 45,444.25 | 929.5 | 9.81 | 245 |
| 1 | 63,386 | 1317 | 16.81 | 416 |

### Dataset preprocessing

The large dimensions and the disproportion between the height and width of the images required the use of patchify process to perform tree ring border segmentation [41,42]. Before each model training session, the increment tree core images assigned to the training and validation were divided into patches. Once the model was calibrated, predictions were made by processing the increment tree core image in patches through a sliding window approach. Previous studies have presented strategies based on small prediction windows (i.e. 128x128 pixels) containing one or two growth tree ring borders [36], or a large prediction window (i.e. 1024x1024 pixels) containing a set of tree ring borders [37]. In both cases, the window sliced the core image along both axes (Fig 2). Thus, the patchify processes had to deal with patch edge regions lacking context, which were compensated by an overlap between windows during core prediction. Here, we performed predictions at a 50% step of the total window, with the average value of the predictions made at each pixel chosen as the pixel probability. Thus, to run only a window slicing along the x-axis and eliminate the uncertainty of information loss in the y-axis, a dataset was created where all the images of the cores were resampled to a specific pixel height while maintaining the aspect ratio (Fig 2).

**Fig 2.**
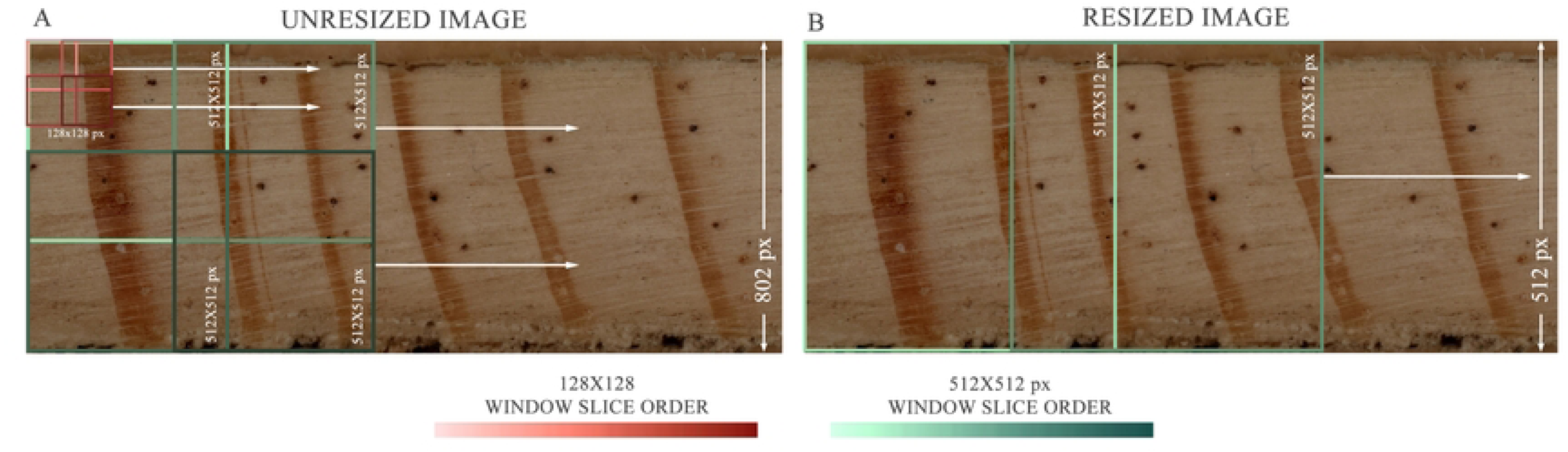
Example of two patchify processes applied to an unresized image and a resized image. Purple boxes represent windows of size 128×128 pixels, and green boxes represent windows of size 512×512 pixels. (a) Display using the original core image with a height of 802 pixels. (b) Display after resizing the core image to a height of 512 pixels while maintaining its aspect ratio.

Multiple preprocessing techniques were applied to the increment tree core images to generate various datasets, which aimed to analyze their interaction with the CNN-based architectures employed and how this interaction was expressed in the segmentation capability of the system.

### Positioning of core growth

The increment tree core images exhibited growth in both directions along the x-axis, posing a challenge for the neural network in a patch-based prediction context. The main difficulty lay in distinguishing the target transition—from latewood to earlywood—from noise, such as the transition from earlywood to latewood. To address this, two distinct datasets were used: (i) one dataset in which the core images included growth in both directions, and (ii) another dataset where all cores were aligned to the right with growth directed to the left, splitting the sample image if necessary.

### Resizing

Resizing was necessary due to the wide range of height and width dimensions presented by the images. Experiments were conducted with those power-of-2 values that fall within the range of image sizes used in previous patchify-based systems [36,37]: 128x128, 256x256, 512x512, and 1024x1024 pixels. As a result, with the first three dimensions, the increment tree core images were downsampled to a greater magnitude as the image size decreases (S1 Figure).

### Image filtering

A Gaussian blur filter with a 15 window size was applied to smooth out cellular details, and a Contrast Limited Adaptive Histogram Equalization (CLAHE) [43,44] filter was used to enhance these details [44,45] with a tile grid size of 8 and a clip value of 40 (S2 Figure).

### Deep learning architectures

#### U-Net Architecture

U-Net is a deep learning neural network architecture initially designed for segmentation in biomedical imaging [25], and extended to different fields due to its good performance [41,46,47]. The architecture consists of two components. The first component is a left-side subnetwork or encoder, which performs down-sampling operations, capturing contextual information and transforming the input image into low-level feature representations across multiple scales [29]. The second is a right-side subnetwork or decoder, which function as the decoding path, carrying out up-sampling operations and mapping the encoder representations back to pixel space through transposed convolutions, facilitating precise localizations [48] (Fig 3). The encoder is composed of blocks, each formed by repeated processes of convolution-batch normalization-Rectified Linear Unit (ReLU) activation. Convolutional layers are used to extract spatial and contextual features from the input tensors, capturing patterns and generating feature maps that help minimize the loss function [49]. Batch Normalization works by normalizing neuron activations within a batch, stabilizing and accelerating training. Finally, ReLU is an activation function that introduces non-linearity into the network, enabling it to learn complex relationships[50]. After applying one or more convolution-normalization-ReLU activation processes, max pooling is used to reduce the spatial resolution of the feature maps (width and height dimensions) while retaining the most relevant information [51]. Feature maps with more abstract and complex characteristics are obtained downwards until the lowest peak, where the way up begins[50]. In decoder, processing blocks are executed with transposed convolution, with the result concatenated through skip connections with the corresponding feature map from the encoder, and convolutional processing of the concatenated feature map[25]. The transposed convolution increases the spatial resolution of the feature map, and the result is concatenated with the corresponding feature map from the encoder to integrate detailed spatial information from that stage of the descending path[25]. Finally, convolutional layers are applied to the concatenated tensor to refine and process the combined features.

**Fig 3.**
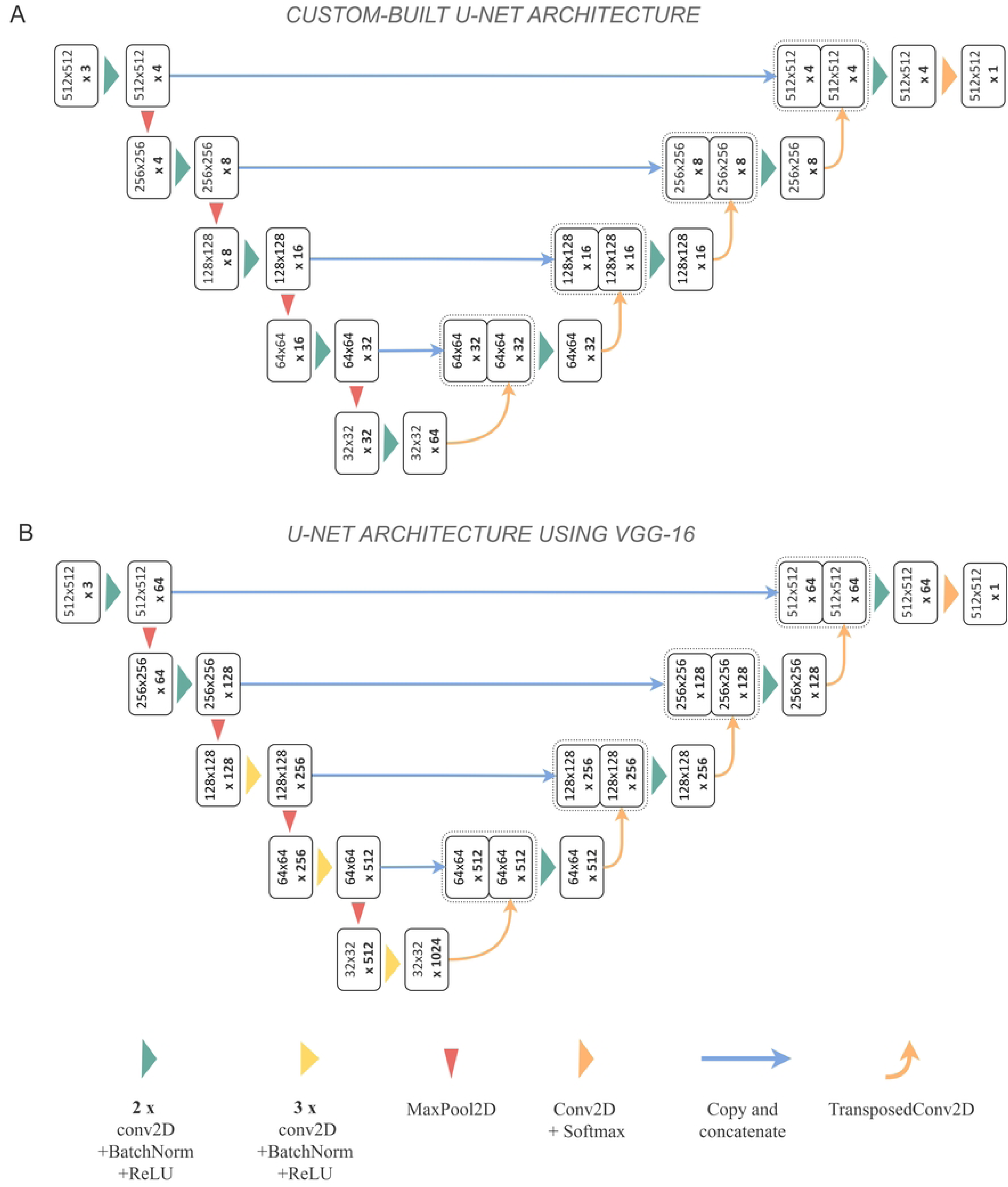
U-Net architectures used in this work. (a) Custom-built U-Net architecture version used in this study and based on [36]. (b) U-Net architecture using VGG-16 as the backbone, employing transfer learning and fine-tuning.

#### Proposed U-Net and variants architectures

Several modifications have been developed based on the original U-Net architecture [25], including Attention U-Net [30] and Attention Residual U-Net [52] architectures, which were chosen in this research for their specific advantages detailed in the descriptions of the custom-built architectures. The design of these custom-built architectures — such as the number of filters per convolutional layer, the number of convolutional blocks at each level, and the total number of levels— followed the guidelines provided by Fabijańska & Cahalan [36]. These researchers applied a progressive reduction in the number of filters per convolutional layer to modify the original Attention U-Net architecture [30], while the performance of the model was monitored until a decrease in capacity was observed compared to the original configuration. This approach reduced the number of parameters and computational cost but also limited the ability to capture complex features and abstract patterns. The custom-built algorithms were initialized with random weights and used a learning rate (lr) 1x10^-3^. Their specific architectures are outlined as follows.

### Custom-built U-Net (CU-Net)

A U-Net architecture (Fig 3) was employed. The convolutional blocks consisted of two 3x3 convolutions, starting with 4 filters in the first block and increasing to 64 in the deepest block, with a 2x2 max pooling applied after each block.

### Custom-built Attention U-Net (CAU-Net)

This architecture was originally proposed by Oktay et al.[30] and incorporated modifications in the skip-connections between the feature maps of the decoder and the encoder. Instead of directly concatenating both layers, an attention mechanism is introduced that weighted the incoming information from the encoder feature maps, deciding which information was relevant and should pass to the decoding stage. Thus, the model learned to suppress irrelevant regions and highlight the important features useful for the segmentation task [30]. In this study, soft attention gates were applied, where the value at each point of the attention map was between 0 and 1, allowing the encoder features to be weighted smoothly.

### Custom-built Attention Residual U-Net (CARU-Net)

Attention U-Net was modified to apply residual convolutional blocks instead of convolutional blocks [52,53]. In this modification, a 1x1 convolution and normalization were applied to the input of the block, which was then connected to the output of the block. Both feature maps were summed, and ReLU activation was applied to the result, so that residual blocks helped to maintain and improve the quality of the feature maps.

#### U-Net using pre-trained models

The custom-built U-Net architectures were optimized to work with image patches containing one or two tree rings [36], whereas the datasets used in this study often include multiple growth tree rings within each patch (Fig 2). We used complex architectures capable of learning more intricate representations and handling data with greater variability (see Table 2). To this end, transfer learning and fine-tuning techniques were used, consisting of adapting previously trained models to address a different task. The employed architecture incorporated the VGG-16 model [54] as encoder subnetwork excluding the dense layers of the original architecture, while the decoder symmetrically reconstructed the encoder subnetwork with randomly initialized weights. Therefore, VGG-16 worked as a feature extractor, since the model had been trained with a large dataset, ImageNet [55], and had learned a good representation of low-level features like spatial, edges, rotation, lighting, shapes [56]. The architecture started at the first convolutional block with 64 filters and reached 1024 at the lowest level and contains convolutional blocks with three convolutional layers (Fig 3).

**Table 2.** Trainable parameters in each architecture.

| Model | Trainable parameters |
| --- | --- |
| CU-Net | 123,763 |
| CAU-Net | 147,165 |
| CARU-Net | 153,537 |
| TLU-Net | 14,097,153 |
| FTU-Net | 21,179,649 |

### U-Net with transfer learning (TLU-Net)

The encoder path had frozen weights throughout the training, while the weights of the decoder were trainable using a 1x10^-3^ lr. This implied that the features previously learned by VGG-16 are sufficient for performing the new tasks.

### U-Net with fine-tuning (FTU-Net)

This approach involved unfreezing a part of the pre-trained model weights for training. Initially, only the decoder was trainable with a 1x10^-3^ lr. From epoch 40 onward, the weights of the last convolutional block of the decoder were also made trainable, with a new lr set to 1x10^-4^. This approach allowed for the adjustment of more complex and abstract features to the dataset used.

### Experimental design

A total of 55 core images, comprising 10,608 tree rings, were used to construct the training, validation, and test sets. Two distinct data partitioning strategies were implemented in the experiments. The first data partitioning strategy, applied only in the first set of experiments, involved training three models with random selection of 40 increment tree cores for training, 10 for validation and 5 for testing. In the second data partitioning strategy, used in the remaining experiments, two rounds of 10-fold cross validation were performed, where in each run, 40 increment tree cores were assigned to train, 10 to validation and 5 to test. The test datasets resulted from the cross-validation processes were referred to as internal test datasets (inTEST). Besides, 6 representative images that captured the diverse challenges addressed, distinct from the 55 core images used for inTest, were reserved for testing outside cross validation. This test dataset was named the external test dataset (exTEST). Finally, the 3 images taken without Macro Flash were referred to as the colored external test dataset (colexTEST). A total of 904 tree rings were contained in exTEST, while 435 tree rings were included in colexTEST. All datasets exhibited tree ring growth toward the left side, except for the experiment described in the fourth set of experiments. Thus, different sets of experiments were conducted. Fig 4 shows that each combination of algorithm and dataset constituted an experiment designed to assess performance under different conditions.

**Fig 4.**
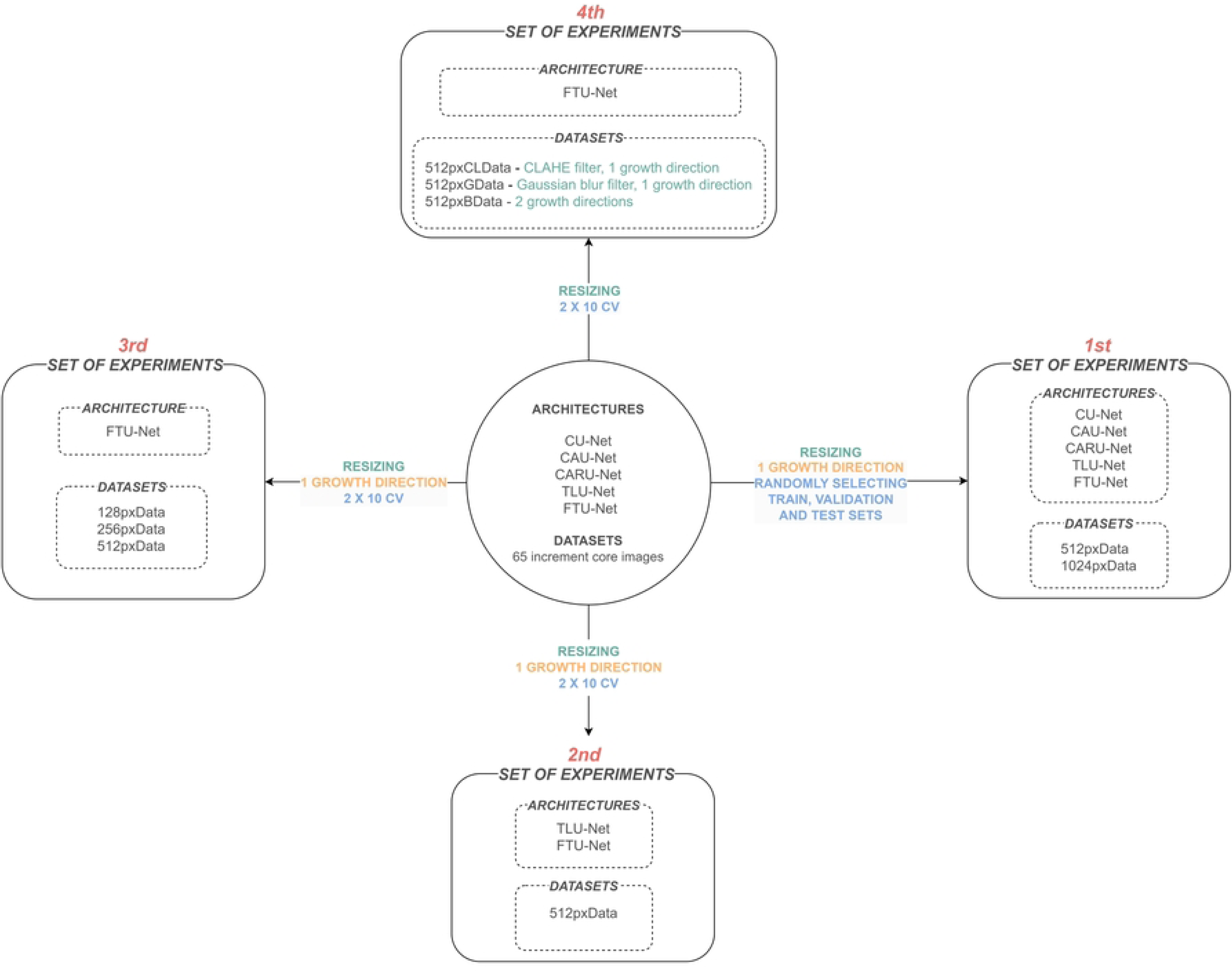
Experimental design scheme. In green, the preprocessing steps performed, and in purple, the partitioning strategies implemented.

### First experiments

The five architectures here considered were trained on two different datasets, where the images were resized to a height of 512 pixels (referred to as the 512pxData) in one case and 1024 pixels (referred to as the 1024pxData) in the other.

### Second experiments

TLU-Net and FTU-Net were executed on 512pxData.

### Third experiments

In these experiments, datasets with image heights of 256 and 128 pixels were generated, referred to as 256pxData and 128pxData, respectively. FTU-Net was executed on datasets 256pxData and 128pxData.

### Fourth experiments

Three datasets were generated from 512pxData: one by applying CLAHE, referred to as 512pxCLData; another by applying a Gaussian blur filter, resulting in 512pxGData; and a third one, exhibiting growth in both directions, referred to as 512pxBData. FTU-Net was executed on 512pxCLData, 512pxGData and 512pxBData.

### Evaluation metrics

An evaluation procedure was applied to analyze the segmentation performance in both normal and abnormal growth patterns across different experiments. In samples with normal growth patterns, the predicted tree ring borders tend to overlap with the ground truth labels. However, achieving precise overlap is less likely in samples with disturbance growth. Previous studies on tree ring segmentation in normal growth patterns have employed an instance-level evaluation [32,34–37], where predicted pixels corresponding to ring borders were grouped into entities and compared with ground truth entities. In contrast, Ge et al. [10] addressed segmentation in abnormal growth patterns and adopted a pixel-level evaluation, which allowed for a more precise assessment of the ability of the models to segment tree ring borders. Here, the evaluation procedure was primarily conducted at the pixel level, complemented by visual inspection. This choice was motivated by the fact that instance-level segmentation required a prior categorization step, which might obscure subtle variations in ring segmentation in abnormal growth patterns. Nevertheless, as a strictly complementary approach, instance-level evaluation was also performed. In pixel-level evaluation, no filtering was applied. However, in instance-level evaluation, a filtering step was introduced, where predicted regions labeled as rings that were shorter than 25% of the image height were removed before evaluation.

To perform a comprehensive comparison between system predictions, we selected the metrics Precision, Recall, Dice score and F1 score to analyze the model results. In the following formulations, true positives (TP) represent the number of positive pixels or instances correctly classified, false positives (FP) refers to the number of negative pixels or instances mistakenly identified as positive, true negatives (TN) denotes the number of negative pixels or instances correctly identified, and false negatives (FN) corresponds to the number of positive pixels or instances wrongly classified as negative. We considered instances correctly classified when the predicted instance and the ground truth had an Intersection over Union (IoU) [57] greater than 0.5; if the IoU was lower, it was classified as a false positive.

### Precision

Measures the proportion of correctly predicted positive pixels or instances (TP) out of all pixels or instances predicted as positive by the model (TP+FP) (equation 1).

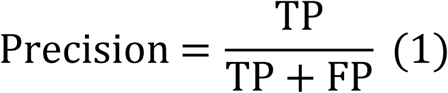

### Recall

Measures the proportion of correctly predicted positive pixels or instances (TP) out of all actual positive pixels or instances (TP+FN) (equation 2).

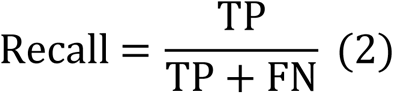

### Dice Similarity Coefficient

Quantify the similarity or the overlap between two sets (equation 3).

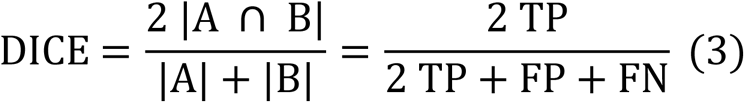

### F1 score

Harmonic mean of precision and recall, balancing false positives and false negatives (equation 4).

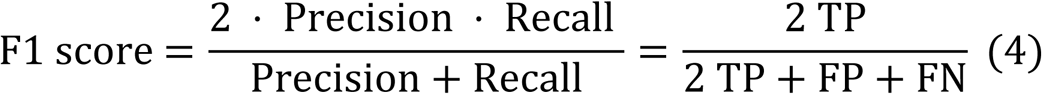

where, in the case of Dice similarity coefficient,|A| is the number of pixels of the predicted mask,|B| is the number of pixels in the ground truth label and |A ∩ B| is number of pixels of the overlapping area. The images used in this study are highly imbalanced. Thus, the use of Dice, which disregards the majority background class, is suitable for evaluating the dataset. As for the F1 score, it was chosen to ensure comparability with previous studies, as this metric is commonly used in tree ring segmentation research. Furthermore, since this is a binary case, the F1 score is mathematically calculated in the same way as the Dice coefficient, and therefore, it also accounts for class imbalance. All the evaluation metrics used in this study range from 0 to 1 and are dimensionless.

### Implementation details

The trainings were conducted on the HPC cluster named Drago, which is part of the Spanish Research Council (CSIC). Two of the nodes are equipped with 512 GB of memory, dual Intel Xeon Gold 6248R processors (24 cores each, running at 3.0 GHz), and 4 NVIDIA Ampere A100 GPUs (80 GB). The storage configuration includes two 240 GB SSD disks and a 10 TB NVMe PCIe 4.0 disk volume. The operating system was Rocky Linux, and the software environment used Python 3.6.8, TensorFlow 2.11, and Keras 2.11. The GPU-based software environment included CUDA 11.7 and cuDNN 8.3.2. Regarding the training hyperparameters, due to class imbalance, the Dice Loss function was used in the training process defined as Dice Loss = 1 ― DICE, the batch size was adjusted according to the available RAM of the GPU used (S1 Table), and all models were trained using the Adam optimizer to adapt the learning rates for each parameter.

## Results

### Similar segmentation performance between resolution maintenance and downscaling

The metric results of the models from the first experiments were grouped into two groups: models trained with the 512pxData and 1024pxData datasets (Table 3). The Dice values of the models trained with 512pxData and 1024pxData were of similar magnitudes, with those trained with 512pxData being slightly higher in inTEST, considerably higher in exTEST, and insignificantly higher in colTEST, showing Dice score differences of 0.017, 0.031, and 0.004, respectively. The models trained with 512pxData required 215 ± 64.4 minutes, while those trained with 1024pxData took three times longer, requiring 684.80 5 ± 241.9 minutes.

**Table 3.** Evaluation pixel metrics resulting from the models of first,second, third and fourth experiments, grouped by datasets and/or architectures.

|  |  |  | Metrics | inTEST | exTEST | colexTEST |
| --- | --- | --- | --- | --- | --- | --- |
| 1ST SET OF EXPERIMENTS | DATASETS | 512pxData | Dice | $0.667 \pm 0.0564$ | $0.653 \pm 0.0822$ | $0.544 \pm 0.1413$ |
| | | | Precision | $0.647 \pm 0.0672$ | $0.696 \pm 0.0567$ | $0.654 \pm 0.0575$ |
| | | | Recall | $0.722 \pm 0.0647$ | $0.656 \pm 0.101$ | $0.512 \pm 0.1741$ |
| | | 1024pxData | Dice | $0.6500 \pm 0.0812$ | $0.622 \pm 0.1298$ | $0.540 \pm 0.1143$ |
| | | | Precision | $0.674 \pm 0.0495$ | $0.715 \pm 0.0551$ | $0.664 \pm 0.0519$ |
| | | | Recall | $0.673 \pm 0.1238$ | $0.600 \pm 0.157$ | $0.492 \pm 0.1535$ |
| | ARCHITECTURES | Custom-built algorithms | Dice | $0.622 \pm 0.0665$ | $0.583 \pm 0.1081$ | $0.466 \pm 0.1089$ |
| | | | Precision | $0.631 \pm 0.0607$ | $0.678 \pm 0.057$ | $0.642 \pm 0.0622$ |
| | | | Recall | $0.665 \pm 0.1151$ | $0.571 \pm 0.1435$ | $0.413 \pm 0.1511$ |
| | | Pre-trained algorithms | Dice | $0.713 \pm 0.0192$ | $0.720 \pm 0.0147$ | $0.655 \pm 0.0165$ |
| | | | Precision | $0.704 \pm 0.0153$ | $0.746 \pm 0.0113$ | $0.684 \pm 0.024$ |
| | | | Recall | $0.747 \pm 0.04$ | $0.715 \pm 0.0317$ | $0.635 \pm 0.0381$ |
| 2ND SET OF EXPERIMENTS | ARCHITECTURES | TLU-Net | Dice | $0.708 \pm 0.0337$ | $0.730 \pm 0.0040$ | $0.672 \pm 0.0087$ |
| | | | Precision | $0.705 \pm 0.0313$ | $0.748 \pm 0.0117$ | $0.688 \pm 0.0205$ |
| | | | Recall | $0.736 \pm 0.0459$ | $0.732 \pm 0.0138$ | $0.662 \pm 0.0278$ |
| | | FTU-Net | Dice | $0.710 \pm 0.0322$ | $0.737 \pm 0.0018$ | $0.677 \pm 0.0081$ |
| | | | Precision | $0.697 \pm 0.0243$ | $0.741 \pm 0.0046$ | $0.691 \pm 0.0135$ |
| | | | Recall | $0.749 \pm 0.0470$ | $0.749 \pm 0.0059$ | $0.668 \pm 0.0248$ |
| 3 RD SET OF EXPERIMENTS | DATASETS | 512pxData | Dice | $0.710 \pm 0.0322$ | $0.737 \pm 0.0018$ | $0.677 \pm 0.0081$ |
| | | | Precision | $0.697 \pm 0.0243$ | $0.741 \pm 0.0046$ | $0.691 \pm 0.0135$ |
| | | | Recall | $0.749 \pm 0.0470$ | $0.749 \pm 0.0059$ | $0.668 \pm 0.0248$ |
| | | 256pxData | Dice | $0.710 \pm 0.0266$ | $0.737 \pm 0.0038$ | $0.698 \pm 0.0056$ |
| | | | Precision | $0.649 \pm 0.0227$ | $0.694 \pm 0.0132$ | $0.663 \pm 0.0117$ |
| | | | Recall | $0.811 \pm 0.0453$ | $0.802 \pm 0.0146$ | $0.74 \pm 0.0151$ |
| | | 128pxData | Dice | $0.660 \pm 0.0317$ | $0.682 \pm 0.0099$ | $0.673 \pm 0.0071$ |
| | | | Precision | $0.559 \pm 0.0399$ | $0.585 \pm 0.0282$ | $0.569 \pm 0.0167$ |
| | | | Recall | $0.837 \pm 0.0649$ | $0.831 \pm 0.0368$ | $0.825 \pm 0.0267$ |
| 4TH SET EXPERIMENTS | DATASETS | 512pxData | Dice | $0.710 \pm 0.0322$ | $0.737 \pm 0.0018$ | $0.677 \pm 0.0081$ |
| | | | Precision | $0.697 \pm 0.0243$ | $0.741 \pm 0.0046$ | $0.691 \pm 0.0135$ |
| | | | Recall | $0.749 \pm 0.0470$ | $0.749 \pm 0.0059$ | $0.668 \pm 0.0248$ |
| | | 512pxGData | Dice | $0.689 \pm 0.0316$ | $0.709 \pm 0.0063$ | $0.624 \pm 0.0236$ |
| | | | Precision | $0.700 \pm 0.0274$ | $0.740 \pm 0.0072$ | $0.731 \pm 0.025$ |
| | | | Recall | $0.710 \pm 0.0459$ | $0.696 \pm 0.0129$ | $0.55 \pm 0.0379$ |
| | | 512pxCLData | Dice | $0.712 \pm 0.0321$ | $0.741 \pm 0.0036$ | $0.693 \pm 0.0046$ |
| | | | Precision | $0.693 \pm 0.0255$ | $0.735 \pm 0.0047$ | $0.671 \pm 0.0091$ |
| | | | Recall | $0.752 \pm 0.0474$ | $0.759 \pm 0.0089$ | $0.717 \pm 0.0158$ |
| | | 512pxBData | Dice | $0.650 \pm 0.0418$ | $0.686 \pm 0.0052$ | $0.537 \pm 0.0181$ |
| | | | Precision | $0.672 \pm 0.0357$ | $0.718 \pm 0.0086$ | $0.588 \pm 0.0257$ |
| | | | Recall | $0.659 \pm 0.0510$ | $0.676 \pm 0.0111$ | $0.5 \pm 0.0260$ |

Furthermore, in inTEST and exTEST, the models trained with 1024pxData showed more variance than those trained with 512pxData. These results indicated that the models trained with both datasets demonstrated comparable segmentation capabilities in terms of metrics, but with a notable difference in computational time.

### Pre-trained models outperform custom-built U-Net models

All custom-built U-Net models demonstrated lower predictive performance compared to the pre-trained models (Table 3). Fig 5 visually illustrates the relationship between predictive performance on exTEST and the computational time required by the different models trained in the first set of experiments. CU-Net, CAU-Net, and CARU-Net demonstrated improvements in Dice scores achieved with 1024pxData compared to 512pxData, albeit at the cost of greater variability in the results. Conversely, pre-trained models performed worse when the resolution of the images was maintained. Only the CAU-Net and CARU-Net models trained with 1024pxData slightly approached the performance of the pre-trained models, although a noticeable gap remained. However, this came at the expense of significantly longer training times. These patterns were also observed in inTEST and colexTEST (S3 Figure).

**Fig 5.**
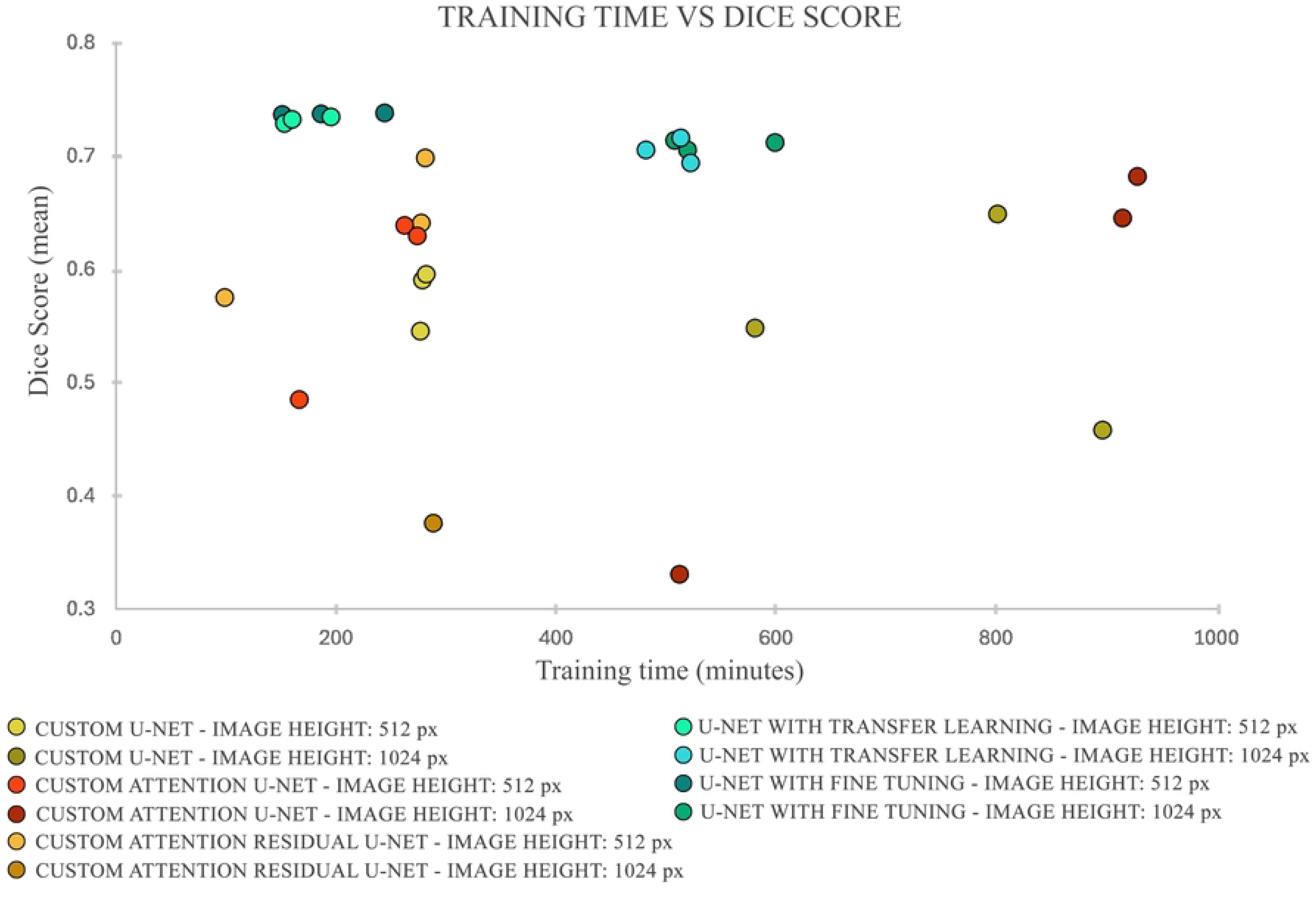
Relationship between the average Dice metric value on exTEST and the training time for first experiments coded by algorithm and dataset.

Thus, pre-trained models demonstrated superior segmentation capability (Dice score = 0.720) compared to the custom-built models (Dice score = 0.583) in exTEST (Table 3). Our analysis suggested that models from the pre-trained group required less training time (352.638 ± 187.354 minutes) compared to the other group (515.151 ± 338.977 minutes). Pre-trained models also showed better performance in both Precision and Recall, with improvements of 0.068 and 0.144, respectively, indicating lower commission and omission errors in exTEST. In inTEST and colexTEST, the pre-trained models also showed notably better performance in Dice metrics, with differences of 0.091 and 0.189, respectively. All these results suggest that pre-trained models can make better predictions than custom-built models.

### Similar performance of U-Net with transfer learning and U-Net with fine-tuning

The metrics obtained from exTEST, colexTEST and inTEST for the results of the second set of experiments were summarized in (Table 3). The paired t-test (p-value < 0.001) supported the hypothesis that TLU-Net (Dice score = 0.730) and FTU-Net (Dice score = 0.737) performed unequal in exTEST, according to the Dice metric. Similarly, the Wilcoxon Signed Rank Test (p-value = 0.007) confirmed a statistically significant difference in colexTEST, where FTU-Net (Dice score = 0.672) achieved a positive difference of 0.005 with respect TLU-Net. However, the Wilcoxon Signed Rank Test (p-value = 0.165) and Levene’s Test (p- value = 0.968; F value = 0.0016) suggested that both algorithms exhibited equality of medians and variances in inTEST. The two algorithms did not exhibit visually discernible patterns of difference in their predictions (S4 Figure). At the instance level, FTU-Net exhibited a negligible improvement of 0.003 and 0.006 over TLU-Net in F1 score on the inTEST and colexTEST datasets, respectively, with no statistically significant differences in central tendency (S2 Appendix). Additionally, according to Levene’s Test, there were no significant differences in variance in either the inTEST dataset (p-value = 0.9897; F value = 0.0002) or colexTEST dataset (p-value = 0.9333; F value = 0.0071), indicating that both models exhibited similar behavior. In the exTEST dataset, FTU-Net showed a minor improvement of 0.008 over TLU-Net in F1 score, which, despite its minor magnitude, was statistically significant (S2 Appendix). Therefore, although some differences in the predictive metrics of the two algorithms were in some test datasets statistically significant, their magnitude was negligible and did not result in perceptible visual differences.

### Visual evaluation of reference model

The FTU-Net models trained on the 512pxData demonstrated competent segmentation performance in detecting tree ring borders within normal growth patterns, generating continuous and separated entities (Fig 6). However, in abnormal growth patterns, specifically when the contrast between latewood and earlywood was very elusive or in the presence of growth disturbance such as callus tissue or compression wood, the model segmented only partial or none (Fig 7). Furthermore, the presence of dark brown stains crossing the growth tree rings horizontally (see example LEFT_PT4_51_C in Fig 7) generally hindered the generation of continuous segmented entities. Nevertheless, when the stain crossed the latewood to a lesser extent and its coloration was lighter, there were more cases where the segmentation achieved continuity. Regarding the sets of narrow tree rings, the models consistently demonstrated the ability to predict continuous and separated tree ring borders up to 200 μ (512 PIXEL-HEIGHT in Fig 8). Beyond this threshold, as the distances decreased, errors became more frequent, particularly near 100 μ, where the segmented tree ring borders eventually merged at some points. As shown in Figure 6, a higher presence of abnormal growth patterns was associated with lower Dice and F1 metrics. The sample RIGHT_PT4_80_C in exTEST deviated significantly from the rest of the group (0.398 Dice and 0.314 F1 scores), making it an atypical value that influenced the mean evaluation metrics, skewing them toward values that reflected poorer overall performance. Nonetheless, the mean was used instead of the median, as it allowed for more precise monitoring of the impact on the worst-performing samples.

**Fig 6.**
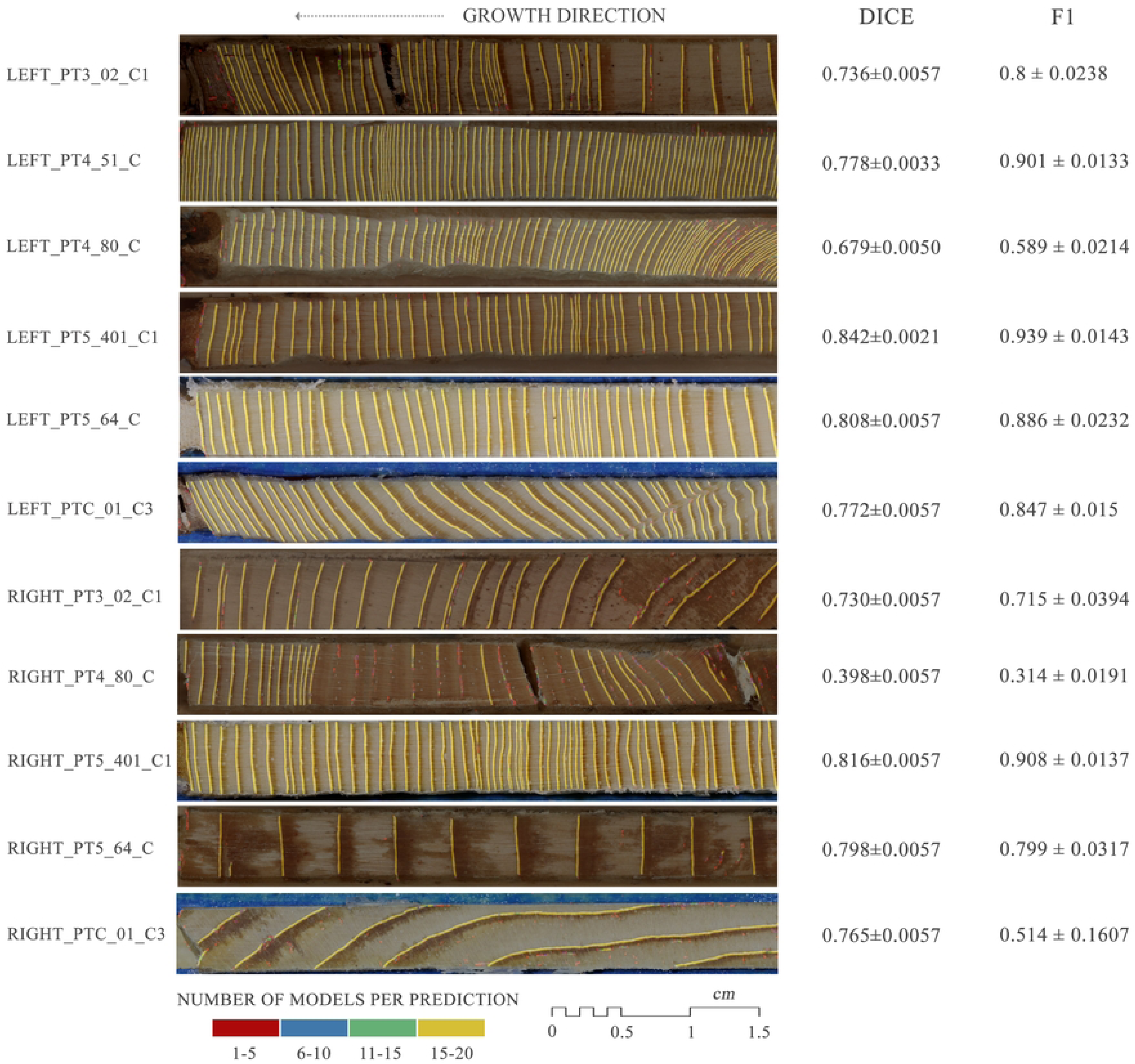
Visual results on tree-ring boundary segmentation and per-image Dice coefficient and F1 score mean and standard deviation for the FTU-Net model across exTEST. The color mapping shows the range of ring border predictions, based on the sum of models.

**Fig 7.**
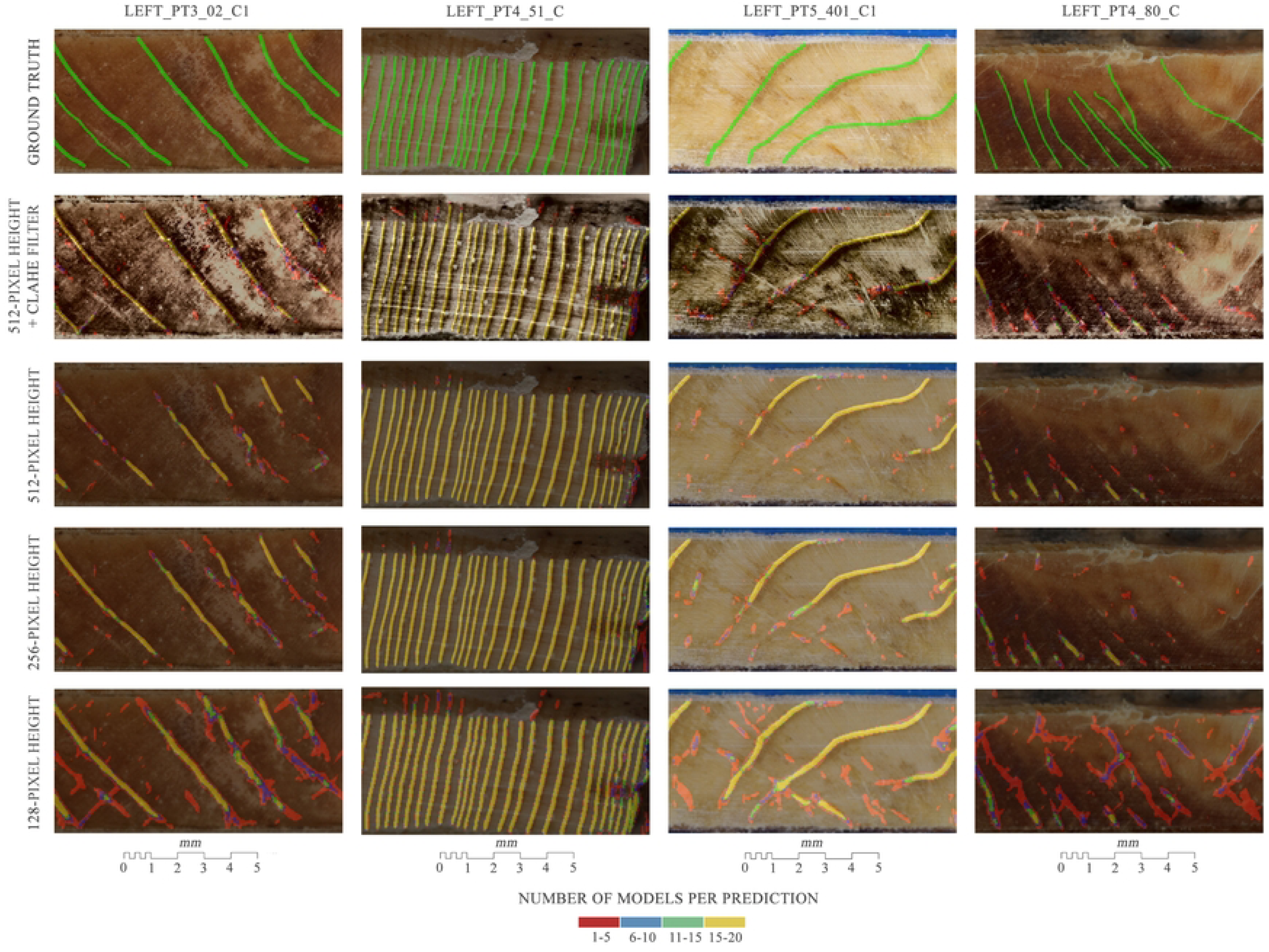
Segmentation of rings in abnormal growth patterns performed by fine-tuned U-Net models trained on datasets with different heights and filters in exTEST. The color mapping shows the range of ring border predictions, based on the sum of models.

**Fig 8.**
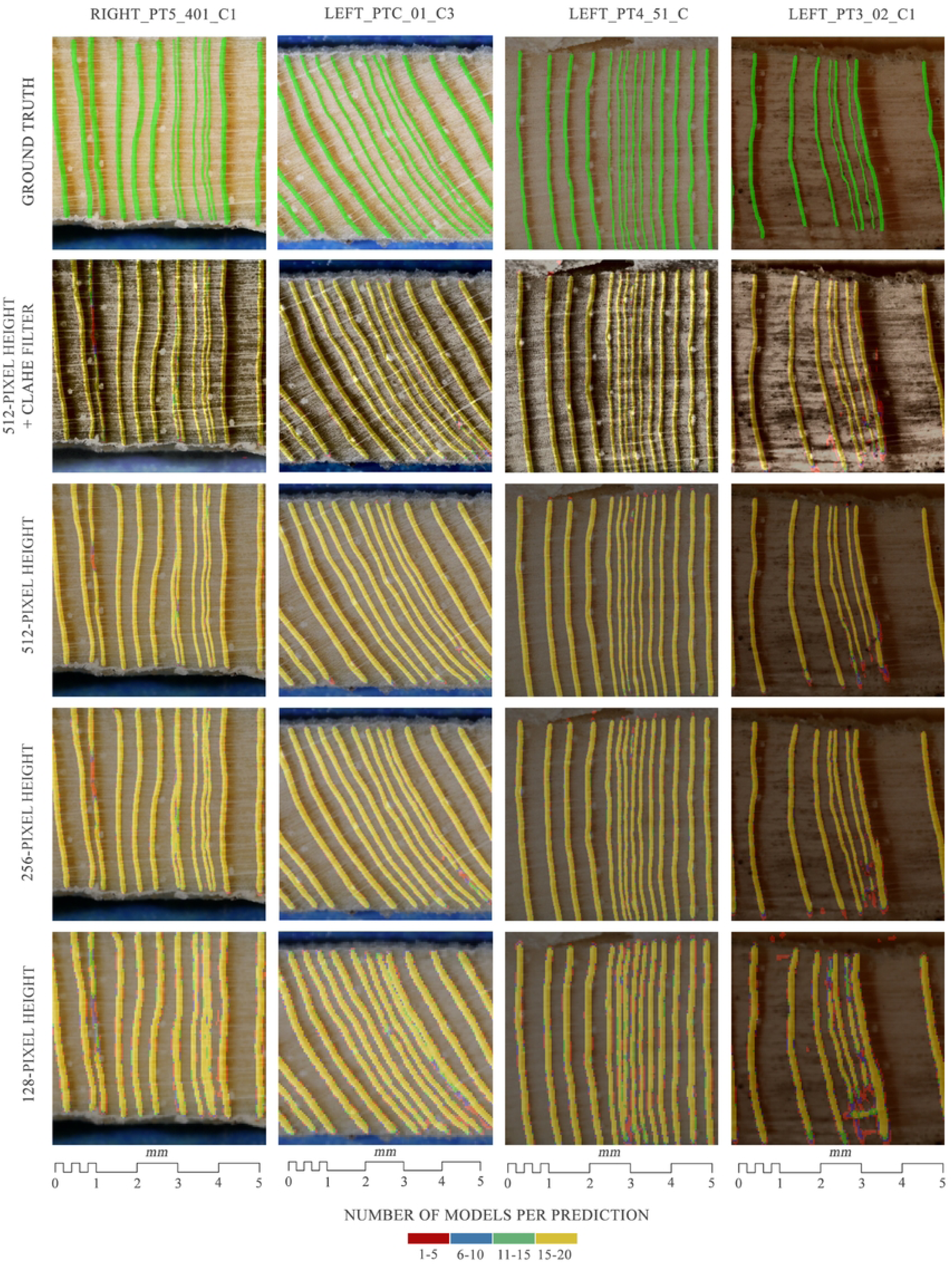
Segmentation of the most closely spaced rings performed by fine-tuned U-Net models trained on datasets with different heights and filters in the external test dataset. The color mapping shows the range of ring border predictions, based on the sum of models.

### Finding model performance limits conditioned by image resolution

In the third set of experiments, metrics were obtained for FTU-Net and trained on datasets with varying height sizes, evaluated on exTEST, colexTEST and inTEST (Table 3). The Dice scores on inTEST, exTEST, and colexTEST for models trained on 512pxData and 256pxData were similar in magnitude, with no change in the first two cases and an increase of 0.021 in the latter when using 256pxData. However, Dice scores dropped for models trained on 128 pxData compared to 256pxData, with decreases of 0.05, 0.055, and 0.025 for inTEST, exTEST, and colexTEST, respectively. A statistical analysis was conducted to assess whether reductions in image size led to significant changes in the Dice Score (Table 4). The Wilcoxon signed-rank exact test applied to inTEST (p-value = 0.8695) and the paired t-test applied to exTEST (p-value = 0.4406) supported the hypothesis that models trained with 512pxData and 256pxData performed equally in terms of central tendencies in Dice values. However, while the Levene’s test indicated homogeneity of variance in inTEST (p-value = 0.7055: F value = 0.145), the F-test applied to exTEST revealed a lack of homogeneity in variances (p-value = 0.0024; F value = 0.23062). The Wilcoxon signed-rank exact test on colexTEST (p-value < 0.001) indicated that the average Dice considerable improvement of 0.021 achieved by reducing the image size was statistically significant. For models trained with 128pxData compared to those trained with 256pxData, all test datasets exhibited different distributions. A Wilcoxon signed rank exact test applied to inTEST (p-value < 0.001), paired t-tests applied to exTEST (p-value < 0.001) and colexTEST (p-value < 0.001) confirmed significant differences in central tendencies across the three datasets. Therefore, the statistical tests demonstrated that the important negative changes observed in 128pxData were also statistically significant. All models trained with the three different test datasets exhibited lower Precision compared to Recall, with differences of 0.008 for 512pxData, 0.108 for 256pxData, and 0.256 for 128pxData in exTEST. The difference between these metrics became more pronounced as the image resolution decreased, due to a reduction in Precision and an increase in Recall.

**Table 4.** Results of the pairwise Dice score comparison of FTU-Net models on test datasets in different resized datasets in third and fourth sets of experiments.

|  | Datasets | Test dataset | Type test | Test statistic (value) | p-value | Significance | Mean Dice score difference<br>(improvement + / worsening - relative to the reference dataset) |
| --- | --- | --- | --- | --- | --- | --- | --- |
| 2ND SET OF EXPERIMENTS | TLU-Net (reference) vs FTU-Net | inTEST | Wilcoxon signed rank exact test | V (67) | 0.165 | Not Significant | +0.02 |
| | | exTEST | Paired t-test | t (-6.7949) | $1.73 \times 10^{-3}$ | Significant (*, p < 0.05) | +0.07 |
| | | colexTEST | Paired t-test | t (-3.0016) | $7.33 \times 10^{-3}$ | Significant (*, p < 0.05) | +0.05 |
| 3RD SET OF EXPERIMENTS | 512pxData (reference) vs 256pxData | inTEST | Wilcoxon signed rank exact test | V (110) | 0.8695 | Not Significant | 0 |
|  |  | exTEST | Paired t-test | t (0.78762) | 0.4406 | Not Significant | 0 |
| | | colexTEST | Wilcoxon signed rank exact test | t (-10.178) | $3.96 \times 10^{-9}$ | Significant (*, p < 0.05) | +0.021 |
| | 256pxData (reference) vs 128pxData | inTEST | Wilcoxon signed rank exact test | V (210) | $1.91 \times 10^{-6}$ | Significant (*, p < 0.05) | -0.05 |
| | | exTEST | Paired t-test | t (22.651) | $3.28 \times 10^{-15}$ | Significant (*, p < 0.05) | -0.055 |
| | | colexTEST | Paired t-test | t (16.191) | $1.43 \times 10^{-12}$ | Significant (*, p < 0.05) | -0.025 |
| 4TH SET OF EXPERIMENTS | 512pxData (reference) Vs 512pxGData | inTEST | Wilcoxon signed rank exact test | V (210) | $1.91 \times 10^{-6}$ | Significant (*, p < 0.05) | -0.021 |
| | | exTEST | Paired t-test | t (19.144) | $7.04 \times 10^{-14}$ | Significant (*, p < 0.05) | -0.028 |
| | | colexTEST | Wilcoxon signed rank exact test | V (210) | $1.91 \times 10^{-6}$ | Significant (*, p < 0.05) | -0.053 |
|  | 512pxData (reference) Vs | inTEST | Wilcoxon signed rank exact test | V (87) | 0.5217 | Not Significant | +0.002 |
| | | exTEST | Paired t-test | t (-3.2773) | $3.96 \times 10^{-3}$ | Significant (*, p < 0.05) | +0.004 |
| | 512pxCLD<br>ata | colexTES<br>T | Wilcoxon signed rank<br>exact test | V (0) | $1.91 \times 10^{-6}$ | Significant<br>(*, $p < 0.05$) | +0.016 |
| | 512pxData<br>(reference)<br>vs<br>512pxBData | inTEST | Wilcoxon signed rank<br>exact test | V (210) | $1.91 \times 10^{-6}$ | Significant<br>(*, $p < 0.05$) | -0.06 |
| | | exTEST | Paired t-test | t (44.337) | $< 2.2 \times 10^{-16}$ | Significant<br>(*, $p < 0.05$) | -0.051 |
| | | colexTES<br>T | Wilcoxon signed rank<br>exact test | V (210) | $1.91 \times 10^{-6}$ | Significant<br>(*, $p < 0.05$) | -0.14 |

**Table 5.** Instance segmentation results and evaluation metrics in tree ring segmentation studies.

|  | N trained models | IoU Overlapped criteria (%) | Precision | Recall | F1 |
| --- | --- | --- | --- | --- | --- |
| Poláček et al. [37] | 1 | 50 | 0.96 | 0.98 | 0.9699 |
| Marichal & Randall [32] | 1 | - | 0.755 | 0.797 | 0.775 |
| Wu et al. [34] | 1 | 25 | 0.6592 | 0.8527 | 0.7348 |
| Kim et al. [35] | 1 | 90 | - | 90.5 | - |
| Fabijańska & Cahalan [36] | 3 | - | 0.99 | 0.92 | 0.9537 |
| FTU-Net with 512pxData | 20 | 50 | 0.852±0.008 | 0.778±0.0067 | 0.813±0.0068 |

At the instance level, using 256pxData instead of 512pxData led to a notable increase of 0.056 in F1 score values for colexTEST, while the differences in exTEST and inTEST were smaller, with unimportant increases of 0.009 and 0.005 in F1 scores, respectively. These improvements were statistically significant in exTEST and colexTEST (S2 Appendix). In contrast, the increasing of F1 score in inTEST was not statistically significant in terms of central tendency, nor did it show a significant difference in variance according to Levene’s test (p-value = 0.6047; F value = 0.2724). Conversely, when comparing 256pxData to 128pxData, the latter showed a substantial decline in F1 score, decreasing by 0.191 in inTEST and 0.170 in exTEST, along with a slight decline of F1 score 0.082 in colexTEST. All three reductions were statistically significant (S2 Appendix). Thus, the changes in the F1 score followed the same trends as the Dice score.

With the reduction of dimensions along the x-axis, the distance between tree rings also decreased, posing a problem for closely spaced tree rings. As shown in Fig 8, models trained with 256pxData showed a decrease in segmentation performance for tree rings with distances between 150 and 200 μm, but maintained consistent performance for distances greater than 200 μm. Finally, models trained with 128pxData exhibited errors up to the boundary of tree rings separated by 300 μm. In S5 Figure, it was observed that a few models (fewer than 5) trained with 256pxData exhibited slightly higher commission errors, over segmenting small areas compared to 512pxData. However, these additional errors were minimal and were offset by slightly lower omission errors, resulting in identical Dice scores for both datasets. In contrast, 128pxData showed an increase in the size of erroneous segmented objects and affected a larger number of models, leading to a marked decrease in Dice scores. Nevertheless, the segmentation remained correct in normal growth patterns and some models (fewer than 5) trained with 128pxData partially segmented tree ring borders in callus tissue where models trained on other datasets failed.

### Invariant performance between raw and filtered imagery

Using the most complex architecture (FTU-Net) and the dataset (512pxData) with the highest resolution, while also being efficient in training time, as the reference system, the fourth set of experiments was used to evaluate the application of different filters to the image data.

Utilizing 512pxData with a Gaussian filter applied (512pxGData) resulted in a statistically significant and substantial deterioration in the Dice score across all test datasets. In both inTEST (p-value < 0.001) and colexTEST (p-value < 0.001), Wilcoxon signed-rank exact tests revealed significant changes in central tendencies, with Dice score reductions of 0.021 and 0.053, respectively. In exTEST, a Paired t-test (p-value < 0.001) indicated a statistically significant deterioration of 0.028 in Dice score. The instance evaluation showed the same trend, with a statistically significant and considerable decrease across all three test datasets (S2 Appendix). Additionally, the segmentation of tree ring borders visually deteriorated in both normal and abnormal growth patterns (S6 Figure).

Regarding the use of CLAHE as a filter (512pxCLData), the paired t-test (p-value = 0.0039) and Wilcoxon signed rank exact test (p-value < 0.001) indicated that the models exhibited different central tendencies in Dice scores on exTEST and colexTEST when 512pxCLData was used. Despite this, the improvement in Dice score was minimal, with an increase of only 0.004 in exTEST and a more noticeable, though still modest, increase of 0.016 in colexTEST. In the inTEST analysis, the Wilcoxon Signed Rank Test (p-value = 0.5217) and Levene’s test (p-value = 0.8495; F value = 0.0365) indicated no statistically significant differences in terms of central tendency and variance in Dice scores. Thus, these results suggest that both datasets exhibited similar performance capabilities. Instance evaluation indicated no changes in central tendencies of the F1 score (S2 Appendix) for either inTEST or exTEST. Levene’s Test for inTEST showed no differences in F1 score variance (p-value = 0.7952: F value = 0.0683), while the F-test for exTEST also revealed no significant difference in F1 score variance (p-value = 0.6836; F value = 1.2087). However, the considerable improvement at the pixel level in colexTEST was reflected by a minor statistically significant improvement of 0.008 in F1 score with the use of 512pxCLData. In Fig 7, CLAHE did not improve the segmentation of tree ring borders in abnormal growth patterns in exTEST.

This demonstrates that the application of 512pxGData worsened the performance of the models, while 512pxCLData showed subtle changes in the metric values. However, it did not lead to any improvement in the segmentation, except in colexTEST.

### Performance declines without only one growth direction

In the fourth set of experiments, 512pxData was also trained without growth normalization. Consequently, the increment tree cores exhibited growth on both sides (512pxBData).

Training the models on 512pxBData resulted in predictions with a noticeably lower Dice score compared to the reference dataset, with Dice score differences of 0.06, 0.051, and 0.14 in inTEST, exTEST, and colexTEST respectively. These differences in Dice scores were also statistically significant across all test datasets (Table 4). Additionally, 512pxBData showed lower Precision and Recall values compared to 512pxData, indicating that this decline in performance was due to an increase in omission and commission errors (Table 3). At the instance level (S2 Appendix), there were statistically significant and quantitatively important decreases of 0.106, 0.138, and 0.267 in the F1 score for inTEST, exTEST, and colexTEST, respectively. Therefore, the performance of the models severely declined when using 512pxBData.

## Discussion

### Effect of U-Net architecture complexity on tree ring boundary segmentation

The first and second sets of experiments were designed to investigate the impact of different U-Net architectures on tree ring segmentation. Using the first set, our results indicated that the custom U-Net models demonstrated lower performance compared to the pre-trained U-Net models. This discrepancy might have been attributed to the design of the custom U-Net architectures, which were optimized for processing small patches of the increment core containing a limited number of tree rings [36]. Consequently, they might have lacked the necessary learning parameters to effectively handle images encompassing the full height of the core, which contained more information and potential sources of noise. Additionally, in this experiment the cellular details were greater than in the study by Fabijańska & Cahalan [36]. However, as shown in Table 3, the custom U-Net models demonstrated a slight improvement when the image resolution was maintained. This observation contradicted the initial assumption that higher-resolution images might have introduced excessive noise for a simple neural network to handle effectively. Additionally, it was felt that the U-Net had failed to take full account of contextual information [58,59]. This could be related to the fact that modifications of U-Net, such as Attention Gate and Residual convolutional blocks, led to an improvement in the Dice metrics (see Fig 5). These two indicators suggested that there was potential to outperform the large architectures currently applied in tree ring segmentation research [10,34,35,37,38], including the pre-trained U-Net used in this research. Higher performance could be achieved by slightly increasing the number of filters in convolutional layers of the architecture proposed by Fabijańska & Cahalan [36] or by incorporating new modifications to that same architecture, such as adding a transformer component [59]. Our results suggested that pre-trained U-Net architecture achieved better results than the simpler architecture proposed by Fabijańska & Cahalan [36]. With the models resulting from second experiments, we could see that the performance of pre-trained U-Net models was intermediate when compared to the range of F1 score values reported in the tree ring segmentation literature (Table 4), which applied Mask R-CNN architectures or improved U-net architectures [10,34,35,37] and exhibited strong segmentation capabilities for regular tree rings, similar to those observed in the literature. In this set of experiments, it was observed that U-Net with transfer learning and U-Net with fine-tuning had similar performance, showing that the adaptation of more complex features to this dataset did not noticeably improve the ability to segment tree ring borders. This could suggest that the proposed pré- trained U-Net architectures reached their maximum performance or that it is necessary to fine-tune simpler features, highlighting the potential significance of more straightforward features in these models and possibly in the task of segmenting tree ring borders.

### Influence of Image Resolution on Segmentation Performance and Growth Direction Inference

Macroscopic studies of tree ring segmentation research revealed a wide range of cellular detail, with cell boundaries clearly visible [37], moderately visible [36], and not visible [10,32,34,35]. Generally, image quality has a positive correlation with the performance of CNNs [60,61]. However, in many machine learning paradigms, reducing the number of inputs or features is often preferred, as it decreases the number of parameters to be optimized and simplifies the model [62,63]. The use of highly detailed images, while potentially beneficial, can increase model complexity, which may necessitate more sophisticated architecture.

Neural networks with a very large number of layers often face the challenge of higher training errors (loss) compared to their shallower counterparts, as the growing number of parameters makes optimization less tractable [53,62]. This is in line with our results derived from the first set of experiments, where pre-trained U-Net models showed worse performance when maintaining the full image resolution (1024pxData) compared to models using downsampled data. In this context, detailed information could be better exploited by architectures with mechanisms such as Attention U-Net, which improve segmentation performance [30,53].

However, increasing image size and architectural complexity comes with trade-offs, such as GPU memory limitations. These constraints often necessitate a reduction in batch size, which can negatively impact gradient calculations for the loss function and, ultimately, model performance [62].

The performance of CNNs, beyond of image resolution, depended on the visual information and specific characteristics of the predicted object [61,62]. For instance, Sabottke & Spieler [62] found that different diagnostic tasks required different optimal image sizes, and Boom et al. [64] did not find the ideal resolution on the highest available. In contrast, Thambawita et al. [65] achieved peak segmentation performance using the highest resolution. The results from the first and third sets of experiments showed that the segmentation system remained robust to image downsampling, with the main impact observed in the segmentation of densely packed tree ring borders. In the fourth set of experiments, applying CLAHE filter enhanced cellular details but did not notably improve the performance of the models. However, better performance of the models was observed in colexTEST, suggesting that this filter could be beneficial if the research involved images with varying illumination conditions. Furthermore, a Gaussian blur filter was applied in this set of experiments, blurring both the cellular details and the gradient between earlywood and latewood, which led to a notable deterioration in model performance, while downsampling removed a substantial amount of information, but preserved contrast between wood types. However, the reduced resolution gradually squeezes sets of narrow tree rings, reducing the ability of the model to correctly predict tree ring borders in these sets, with the minimum consistent detectable distance increasing from 200 μm to 300 μm. The generally maintained segmentation capacity despite downsampling could imply that CNNs primarily focus on color gradients rather than complex cellular features, suggesting that at the macro level, it might not be necessary to capture cellular detail levels to segment tree rings spaced approximately 300 μm.

The fourth experiment was designed to test if the cellular resolution and tree ring convexity could efficiently detect the growth direction of an increment core. Previous studies [10,32,33] have successfully segmented tree rings in tree cross section, where the direction of growth is multidirectional. In contrast, our models produced significantly worse results when growth direction information for increment cores was not provided beforehand. We believe that the model of Ge et al. [10] achieved superior results because it used the central position of the pith as a reference for growth direction. This idea is supported by other studies [32,33], where one of the tasks of one of the CNNs in their intelligent system is to detect the pith of cross section. In our system, which relied on patches, the convexity of tree rings and the cell details were insufficient. As a result, transitions of colors like random dark wood stains in earlywood and transitions from earlywood to latewood were often misinterpreted as tree ring borders, leading to an increased number of false positives.

### Performance of CNN-Based System in abnormal Growth Patterns

Except for Ge et al.[10], previous studies did not focused on tree-ring segmentation with growth anomalies. Here we worked with samples with very narrow rings, which posed a significant challenge due to their proximity. With 512pxData, we found consistent predictions up to a distance of 200 μm. Below this threshold, the issues progressively increase until 100 μm, where no accurate predictions are achieved. These issues were previously reported in other studies [35–37], where different segmentation instances merged due to proximity. Since they did not provide quantitative distance measures, we are unable to quantitively compare their capabilities. These studies [35,37] used the Mask R-CNN architecture, which is not specialized to fine pixel segmentation as U-Net architectures, and Fabijańska & Cahalan [36] had to merge small patch predictions in both directions due to these challenging situations.

Wu et al.[34] did not study sets of narrow tree rings packages, while Ge et al. [10] claimed to achieve separate segmentation, but their figures showed incorrectly merged tree ring instances, likely due to the small image size used. The system proposed here, based on U-Net and a sliding window with movement along a single axis, consistently segmented tree rings with distances above 200 μm, correctly identifying most of the closely spaced rings in this dataset. Thus, regarding growth disturbances, it can be stated that the proposed system also works correctly in growth suppression, except in some extreme cases with very closely spaced rings. Additionally, the system performs well in detecting growth release, as it did not pose a technical challenge. Following the analysis of growth disturbances observed throughout our experiments, we found that tree rings in reaction wood and in the pith of the increment core were segmented correctly in several instances. However, the models also made several mistakes, which prevented them from being fully functional. Segmentation errors were particularly prevalent in callus tissue, where the performance of the models was significantly worse. Previous studies [32,34–36] reported issues with abnormal growth patterns or directly avoided labeling those tree rings borders, while Ge et al. [10] claimed success, although concerns about segmentation in abnormal growth patterns may arise when inspecting their figures. Our study contributes to improving the performance of segmenting tree ring borders in growth suppression and enhances the understanding that the problem of segmenting tree rings in abnormal growth patterns, such as reaction wood or callus tissue, cannot be solved by using increment tree cores from trees affected by growth disturbances. Different studies work with different species that present varying textures and coloration. Yet CNNs still struggle to adapt in cases where there is no clear color gradient. We believe that a successful experiment to segment tree rings in abnormal growth patterns could involve training CNNs exclusively with the sections of increment cores affected by growth disturbances, allowing the gradients of the network to focus on minimizing this specific problem.

## Conclusions

This work provides valuable insights into the field of tree ring segmentation, particularly in the context of growth disturbance trees, by studying different deep-learning complex algorithms and datasets with varying levels of cellular detail. We found that at a macro level, even with complex neural networks and high-resolution images containing detailed cellular structures, U-Net primarily focus on the simple feature of color gradients. Therefore, U-Net were unable to fully exploit the complex features present in the high-resolution images, suggesting that there is room for improvement in training a neural network capable of capturing these features using U-Net modifications. Our findings show that segmentation, at least with U-Net architectures, remains resilient to image downsampling as long as the color gradient between tree rings remains clear. However, it becomes sensitive in densely packed tree rings, where the capacity of the model to segment consistently decreases from 200 μm to 300 μm. This could be valuable for researchers who are not focused on densely packed tree rings, as they may save time by using lower-resolution images and reallocating resources to other tasks. Another insight is that, when working with increment cores and using the patchify process, even with the highest resolution presented in this work, it was not possible to infer the growth direction without erroneously segmenting the transition from earlywood to latewood as the tree ring border. This underscores the need for incorporating prior knowledge into the system, such as aligning the increment core with the same growth direction. Finally, regarding growth disturbances, we demonstrated that our system robustly segments tree ring borders in cases of growth suppression and growth release. However, for other types of growth disturbances, such as callus tissue and reaction wood, working solely with datasets containing increment cores affected by these disturbances is insufficient to successfully segment tree ring borders, making it necessary to explore alternative systems. Nevertheless, our models achieved some success in specific cases, indicating that there is still potential for further improvements

## Acknowledgment

This work was supported by the EXTreeM project (PID2021-1245730A-100) by the Ministerio de Ciencia e Innovación MCIN/AEI/10.13039/501100011033. We thanks the Forest Planning and Management Service from Government of Aragon for giving us the permission for sampling as well as Gerardo Benito for his support during this study.

## Supporting information

**S1 Figure. Example of LEFT_PT5_401_C1 with different height resized.** (a) 128-pixel height resized image. (b) 256-pixel height resized image (c) 512-pixel height resized image (d) 1024-pixel height resized image (e) 834-pixel height original image.

**S2 Figure. Example of LEFT_PT4_02_C1 with different filters applied**. (a) CLAHE filter. (b) Blur filter. (c) Raw image.

**S3 Figure. Relationship between the average Dice metric value on test sets and the training time for the first set of experiments, coded by algorithm and dataset**. (a) On inTEST. (b) On exTEST. (c) On colexTEST.

**S4 Figure. Predictions on the images from exTEST made by the models from the second set of experiments.** In descending order: image with ground trith labeling, image with TLU- Net prediction, image with FTU-Net prediction.

**S5 Figure. Predictions on the images from exTEST made by the models from the third set of experiments**. In descending order: image with ground truth labeling, image with predictions from the models trained with 512pxData, image with predictions from the models trained with 256pxData, image with predictions from the models trained with 128pxData.

**S6 Figure. Predictions on the images from exTEST by FTU-Net from the fourth set of experiments**. In descending order: image with ground truth labeling, image with predictions from the models trained with 512pxData, image with predictions from the models trained with 512pxGData, image with predictions from the models trained with 512pxCLData, image with predictions from the models trained wth 512pxBData.

**S1 Table. Summary table of the batch size values adjusted for each algorithm-dataset combination.**

**S1 Appendix. Graphic information about the fieldwork trip**. (a) Zone map showing the sampled trees, including the 64 used for tree ring segmentation. (b) Sampled tree with a significant ramification near the soil. (c) Sampled tree with a scar on the trunk. (d) Sampled tree with trunk curvature.

**S2 Appendix. Tables of instance evaluation metrics**. Table 1, evaluation instance metrics resulting from the models of first, second, third and fourth set of experiments, grouped by datasets and/or algorithms. Table 2, results of the pairwise F1 score comparison of experiments.

